# Reformer: Deep learning model for characterizing protein-RNA interactions from sequence at single-base resolution

**DOI:** 10.1101/2024.01.14.575540

**Authors:** Xilin Shen, Xiangchun Li

## Abstract

Protein-RNA interactions play an essential role in the regulation of transcription, translation, and metabolism of cellular RNA. Here, we develop Reformer, a deep learning model that predicts protein-RNA binding affinity purely from sequence. We developed Reformer with 155 RNA binding protein (RBP) targets from 3 cell lines. Reformer achieved high prediction accuracy at single-base resolution when tasking with inferring protein- and cell-type-specific binding affinity. We conducted electrophoretic mobility shift assays to validate high-impact RNA regulation mutations predicted by Reformer. In addition, Reformer learned to capture protein binding motifs that cannot be discovered by eCLIP-seq experiments. Furthermore, we demonstrated that motif signatures related to RNA processing functions are encoded within Reformer. In conclusion, Reformer will facilitate interpretation of the regulation mechanisms underlying RNA processing.

## Background

Critical cellular processes, such as gene expression and proteogenesis, rely on RNA binding protein (RBP) interacting with their target RNAs. RBPs are involved in regulating numerous aspects of RNA processing including RNA splicing, stability, localization, editing, and translation [1]. Defects in these processes are associated with human genetic diseases such as autoimmunity, neuropathic disease, and cancer [2–5]. Hence, characterizing protein-RNA binding from RNA sequences holds promise for a better understanding of the mechanism of RBP binding underlying RNA dysregulation and diseases.

Deep learning methods offer an effective data-centric approach for characterizing protein-RNA interactions. For example, DeepBind predicted the binding affinity of protein-RNA interactions from RNA sequences with a deep convolutional network [6]. RNAProt used a recurrent neural network for predicting protein-RNA binding [7]. PrismNet applied a residual network to integrate RNA sequence and structure information for predicting RBP binding patterns [8]. These models treated RNA-protein interactions as a binary classification task to distinguish binding sites from non-binding sites. None of these methods can be applied for the prediction of RBP-RNA binding at single-base resolution. Besides, these models only considered binding regions of approximately 100 bp [6–8] but disregarded the influence of contextual information surrounding the peaks, which has been shown to impact RBP binding [9].

In this study, we introduce Reformer which is based on transformer [10] aiming to improve prediction resolution and facilitate greater information flow between peaks and their surrounding contexts. We frame the model to predict RBP binding affinity purely from sequence. Training on a large number of RNA sequences across 155 RBP targets from 3 cell lines, we demonstrated a high correlation between the predicted peaks and actual peaks derived from eCLIP-seq. Reformer captured diverse RBP binding patterns in peak regions as well as their surrounding contexts. Using binding information of base resolution, the model can detect deleterious mutations that disrupt RNA-protein interactions that are supported with experimental validations. Reformer provides a unified framework for characterizing RBP binding and prioritizing mutations that affect RNA regulation at base resolution.

## Results

### An overview of Reformer

We developed a deep learning model named Reformer (RNA-protein binding modeling with transformer) to quantitatively characterize RNA-protein binding affinity (**Figure 1**). Reformer was motivated by the bidirectional encoder model [11]. Reformer was designed to predict the binding affinity of RNA-protein interactions from cDNA sequence at single-base resolution. Reformer consists of 12 transformer blocks. Each block had 12 attention heads with 768 hidden units. We developed Reformer on a dataset consisting of 872,618 peaks from 225 eCLIP-seq experiments (**Figure 1A**, see **Methods**) encompassing 155 RNA binding proteins (RBPs) across 3 cell lines. These RBPs span different RNA regulatory functions including RNA export, stabilization and decay [1]. We unified individual peak into 511 bp and tokenized it into a sequence with 3-mer representation. The corresponding eCLIP-seq target of the sequence was added at the beginning of the sequence (**Figure 1B**). For each base, the transformer layer computed a weighted sum across the representations of all other bases of the sequence. This allows Reformer to refine predictions by incorporating information from relevant regions across the entire sequence (**Figure 1C**). Employing a regression layer for coverage prediction, Reformer outputs binding affinities for all bases (**Figure 1B**). The model was trained to minimize the discrepancy between the predicted and actual binding affinity (see **Methods**).

**Figure 1.**
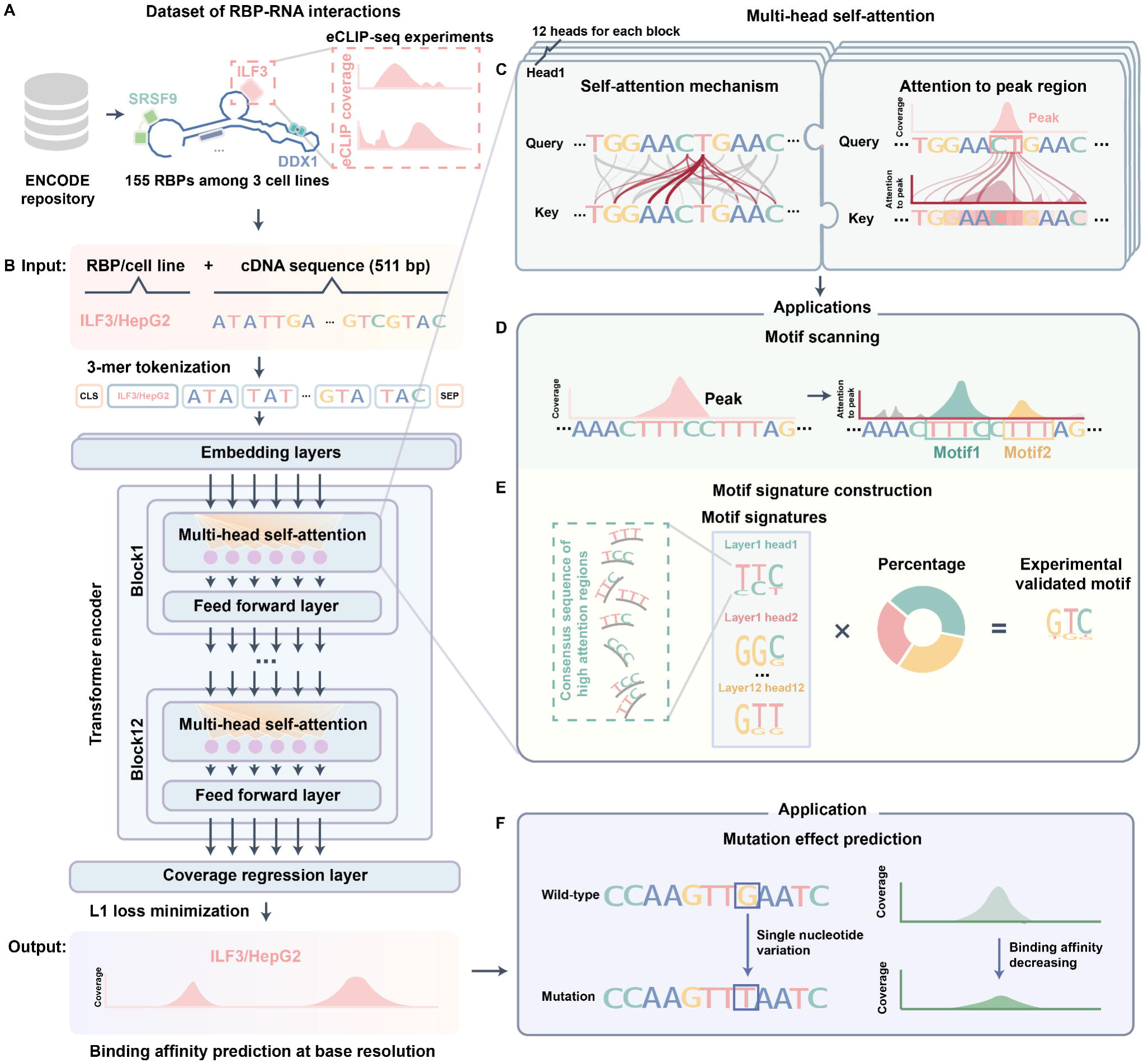
Overview of Reformer. (**A**) Reformer was trained to predict protein-RNA binding affinity for eCLIP-seq experiments. (**B**) The architecture of Reformer was built upon a bidirectional encoder. The model was designed to predict binding affinity between RNA and protein, with cDNA sequence as inputs. (**C**) The self-attention mechanism of Reformer enables the calculation of attentions of individual bases to peak region. (**D-E**) The downstream applications of the self-attention mechanism, include motif enrichment analysis and motif signature compilation. (**F**) Application of the trained Reformer model in predicting the effect of mutations on RBP binding affinity.

We showcase the trained Reformer for three applications: (1) motif enrichment analysis conducted from regions with high attention scores identified by Reformer (**Figure 1D**); (2) compilation of motif signatures with highly attended regions from attention heads, enabling the construction of canonical motifs (**Figure 1E**); and (3) prediction of the effects of mutations on RNA-RBP binding via assessing changes in predicted binding affinity. Our framework will offer insights into the regulatory mechanisms of RNA-RBP interactions and facilitate the fine-mapping of human diseases that are associated with RBP regulation.

### Reformer accurately predicts binding affinity of RNA-RBP interaction at base resolution

We further evaluated the utility of Reformer for RBP-RNA binding prediction. At base resolution, Reformer achieved a Pearson correlation of 0.85 between the predicted and actual affinity (**Figure 2A**). For individual sequences, Reformer yielded high precision in prediction (mean Pearson *r* = 0.70), indicating that the model captured the variation of binding affinity in same sequence (**Figure 2B**). For each sequence, the predicted coverage of the genomic track was more similar to the observed coverage of the same target as compared to other targets (paired t-test, *p*-value < 3e-12; **Supplementary Figure 2**). The difference in peak widths and binding strength can also be captured by Reformer (**Figure 2C**). Reformer can accurately predict the total binding affinity for peaks both across all eCLIP-seq experiments and for individual experiments (mean Pearson *r* = 0.85, 0.60, respectively; **Figure 2D-E**). The similarity between predicted and observed peaks resembles biological replicates (**Supplementary Figure 3**). The difference between predicted coverage from Reformer and actual coverage was 0.61, which is close to the coverage differences between biological repeats (**Figure 2F-G**; paired t-test, *p*-value = 0.21). The difference in coverage between biological repeats was 0.40. The high similarity between predicted and observed peaks was also qualitatively evident (**Figure 2H**).

**Figure 2.**
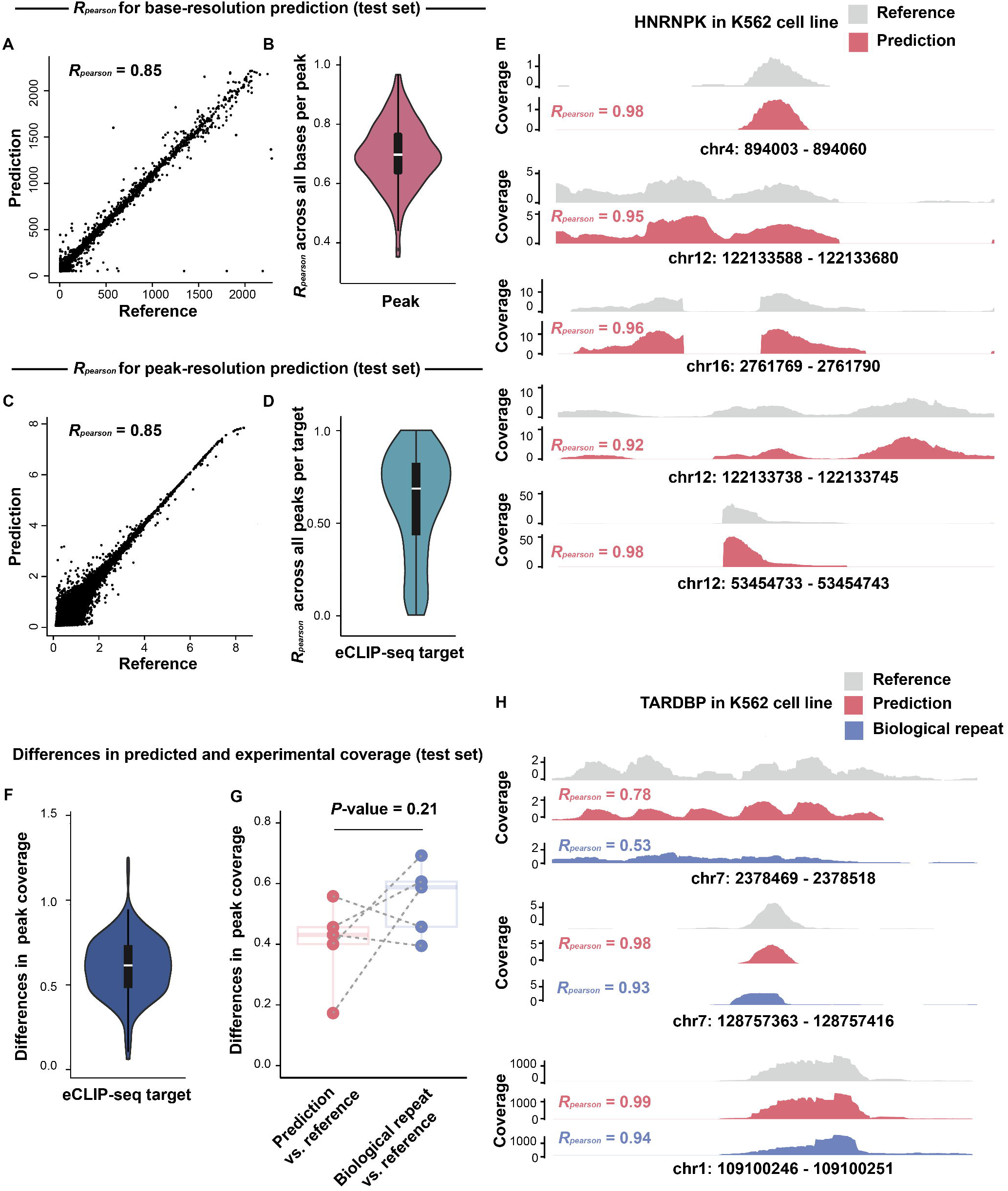
Evaluation of the performance of Reformer in test set. (**A-B**) Pearson correlation between the reference and predicted base-level binding affinity: (**A**) calculated across all sequences, and (**B**) calculated for each individual sequence. (**C-D**) Pearson correlation between the reference and predicted peaks binding affinity: (**C**) calculated across all eCLIP-seq experiments and (**D**) calculated for each individual experiment. (**E**) The examples of reference and predicted genomic tracks of HNRNPK in K562 cell line. (**F**) Violin and boxplot demonstrating the coverage differences between reference and predicted peaks across eCLIP-seq experiments. The boxplot presents the median, upper, and lower quartile of the differences. (**G**) Dot and boxplot illustrating the coverage difference between predicted and reference peaks. The *p*-value was calculated using a paired t-test. (**H**) The examples of reference, predicted and biological replicated genomic tracks of TARDBP in K562 cell line.

### Reformer attends to RBP binding motifs

The attention mechanism underlying transformer architecture [10] can be applied for sequence element discovery [12]. We extracted regions that are most attended by Reformer for each target and inspected whether canonical motifs receive higher attentions in these regions (see **Methods**). We observed that 872 of the 960 validated motifs were significantly enriched in high attention regions (**Figure 3A-B**, *p*-value < 0.05). In comparison, 486 motifs were enriched in peak regions (**Figure 3A-B**, *p*-value < 0.05). A total of 392 motifs can be discovered by Reformer but not in peak regions (**Figure 3C**). These newly identified motifs locate around the binding peaks (**Figure 3D and Supplementary Figure 4**). For example, as a canonical motif of U2AF2 target, TTTTT locates at the left side of eCLIP-seq peaks (**Figure 3D**). ACAA motif for HNRNPL locates at the right side of the peaks and GGATTC motif for HNRNPC locates in the middle of two peaks (**Supplementary Figure 4**).

**Figure 3.**
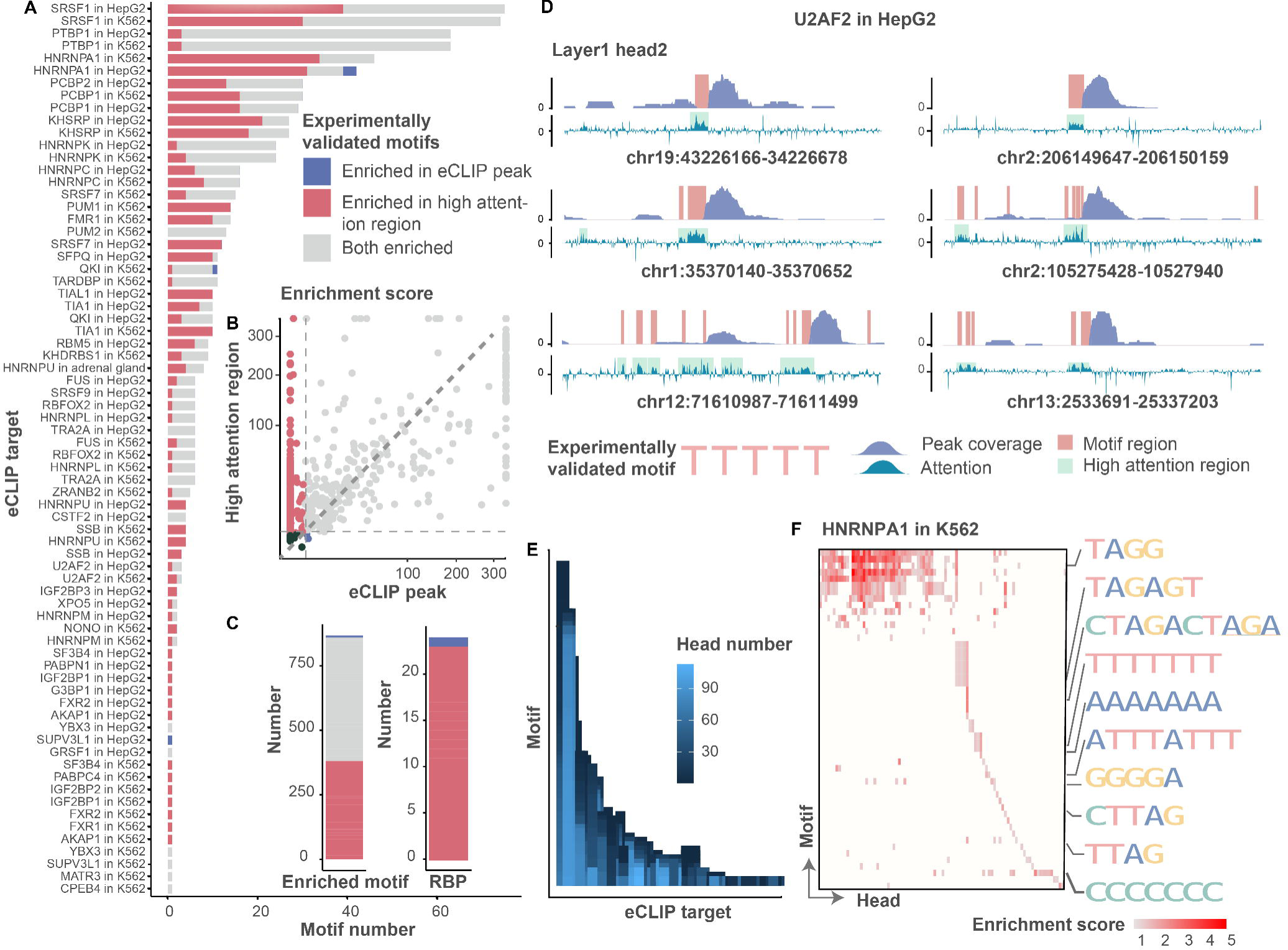
Enrichment of canonical motifs in regions highly attended by Reformer. (**A**) Stacked bar plot showing the target-specific motifs that enriched in eCLIP-seq peaks and Reformer’s high attention regions. Motifs enriched only in eCLIP peaks are colored blue, motifs enriched only in Reformer’s highly attented regions are colored red, and motifs enriched in both regions are colored grey. (**B**) Scatter plot demonstrating the enrichment score of motifs in eCLIP peaks versus high attention regions. Each dot represents a motif. The enrichment score is represented as -log_10_(*p*-value) calculated with AME algorithm. (**C**) Left, the total number of motifs that enriched exclusively in eCLIP peaks, high attention regions, and both regions. Right, corresponding RBPs for motifs exclusively enriched in eCLIP-seq peaks (blue) and high attention regions (red). (**D**) An example describing the enrichment of U2AF2 motifs in genome tracks (HepG2). The red frame represents the peak region validated by eCLIP-seq experiment, while the blue frame represents the high attention regions of layer-1 head-2 of Reformer. (**E**) Bar plot depicting the number of heads in which motifs are enriched. Each column represents an eCLIP-seq target and the stack bar represents a corresponding motif of the target. The color intensity of the bars represents the number of enriched head for each motifs. (**F**) Example highlighting the enrichment of different motifs of HNRNPA1 in different heads.

Most of the motifs (810/872, 93%) were encoded in multiple heads, while 7% of motifs (*n* = 62) were recognized by unique heads (**Figure 3E**). Meanwhile, different heads were enriched for different motifs (**Figure 3E-F, Supplementary Figure 5**). These results demonstrated that Reformer can robustly recognize motifs both in peak regions and their surrounding contexts.

### Deciphering motif signatures from regions highly attended by Reformer

We compiled motif signatures from Reformer. Specifically, we extracted cDNA sequences from regions highly attended by Reformer and subsequently compiled them into one consensus sequence per head (see **Methods**). Each consensus sequence was considered as a motif signature. Eventually, we obtained 78 motif signatures after excluding replicates (**Supplementary Figure 6**). Most of these signatures (n = 77) were significantly different from the consensus sequences constructed from random regions (*p*-value < 0.05; **Supplementary Table 3**). One motif signature TTTTT that achieved marginal statistical significance (*p*-value = 0.06) is a validated motif of multiple RBPs [31–33]. The result indicates that these sequences represented the specific sequence patterns captured by attention heads.

We used NMF2D algorithm [19] to reconstruct 1312 canonical motifs against these 78 motif signatures. 1038 reconstructed motifs achieved significant similarity with the canonical motifs (TOMTOM *q*-value < 0.05; **Supplementary Table 3**). These motifs can be reconstructed with particular motif signatures at different intensities. For example, GCCAA motif can be constructed with Signatures 10 and 75 (**Figure 4B**). Both AAAGG motif of TLR3, TAATT motif of A1CF and GGGG motif of HNRNPH1 can be constructed with Signatures 18, 42, 61 and 77, but with different intensities (**Figure 4C**).

**Figure 4.**
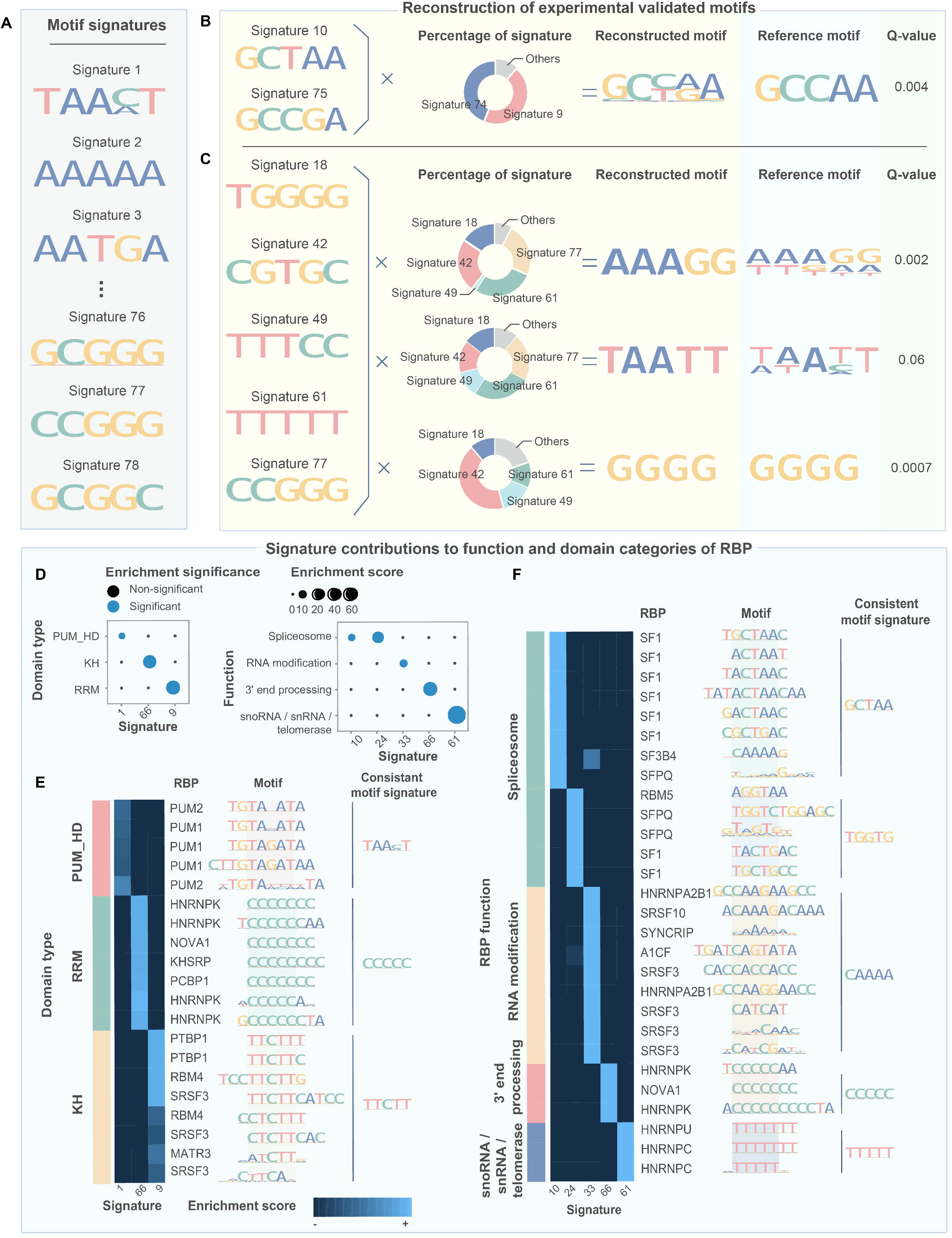
Validation of motif signatures and exploration of their relevance to RNA regulation. (**A**) The motif signatures constructed with high attention regions. (**B**) The examples describing the reconstruction of known motifs using motif signatures of varying intensities. The similarity between the known motif and reconstructed motif was determined using the TOMTOM algorithm, with a q-value < 0.05 considered significant. (**C**) Dot plot illustrating the functional and domain specificity of the motif signatures. The circle size reflects the average contribution score of the signature to RBPs with the same function, and the color reflects whether a signature is significantly enriched in a specific functional category. Significance was determined using the Wilcoxon rank-sum test, with a *p*-value < 0.05 considered significant. (**D-E**) Heatmap representing the contribution score of motif signatures to motifs with specific functions.

We investigated whether the signature contributions are related to the function and domain categories of RBPs encompassed by the motif (see **Methods**). We found that Signatures 1, 66 and 9 were significantly associated with RBPs possessing PUM, KH and RRM domains, respectively (**Figure 4D**, Wilcoxon rank-sum test, *p*-value < 0.05). Notably, Signature 1 (i.e. TAACT) exhibited a strong attribution to the motifs of PUM1 and PUM2 (**Figure 4E**), which has been previously validated in independent research [13]. Signatures 10, 24, 33, 66 and 61 displayed high attribution scores to motifs associated with spliceosome, RNA modification, 3’ end processing and telomerase binding (**Figure 4D**, Wilcoxon rank sum test, *p*-value < 0.05). Visually, RBPs from the same functional categories demonstrated similar motif patterns corresponding to their respective motif signatures (**Figure 4F**). Taken together, these findings suggest a relationship between the motif signatures deciphered by Reformer and RBP functional categories.

### Reformer provides informative predictions about RNA mutation effect on RBP binding

A central application of Reformer was to predict the influence of genetic variations on RBP regulation. We applied Reformer to predict mutation effect on RNA-protein interactions for 553,803 single nucleotide variants (SNVs) collected from TCGA repository, ClinVar database and the 1000 genomes Project (see **Method**) [14,15]. Variations with high scores were pathogenic or related to RNA splicing (**Figure 5A**, **Supplementary Table 5**). For SNVs curated by experts, we observed that pathogenic mutations have significantly higher mutation effect scores than benign mutations (**Figure 5B**; t-test, *p*-value < 1e-5). Mutations proximate to splice sites have higher mutation score (**Supplementary Figure 7**). Quantitatively, mutations located at splice sites have significantly higher mutation scores as compared with missense mutations (**Figure 5C**; t-test, *p*-values < 1e-2). Among splice site mutations, rare mutations had significantly lower mutational effect compared to common mutations (**Supplementary Figure 8**; t-test, *p*-values = 4e-3), suggesting their susceptibility to strong negative selection.

**Figure 5.**
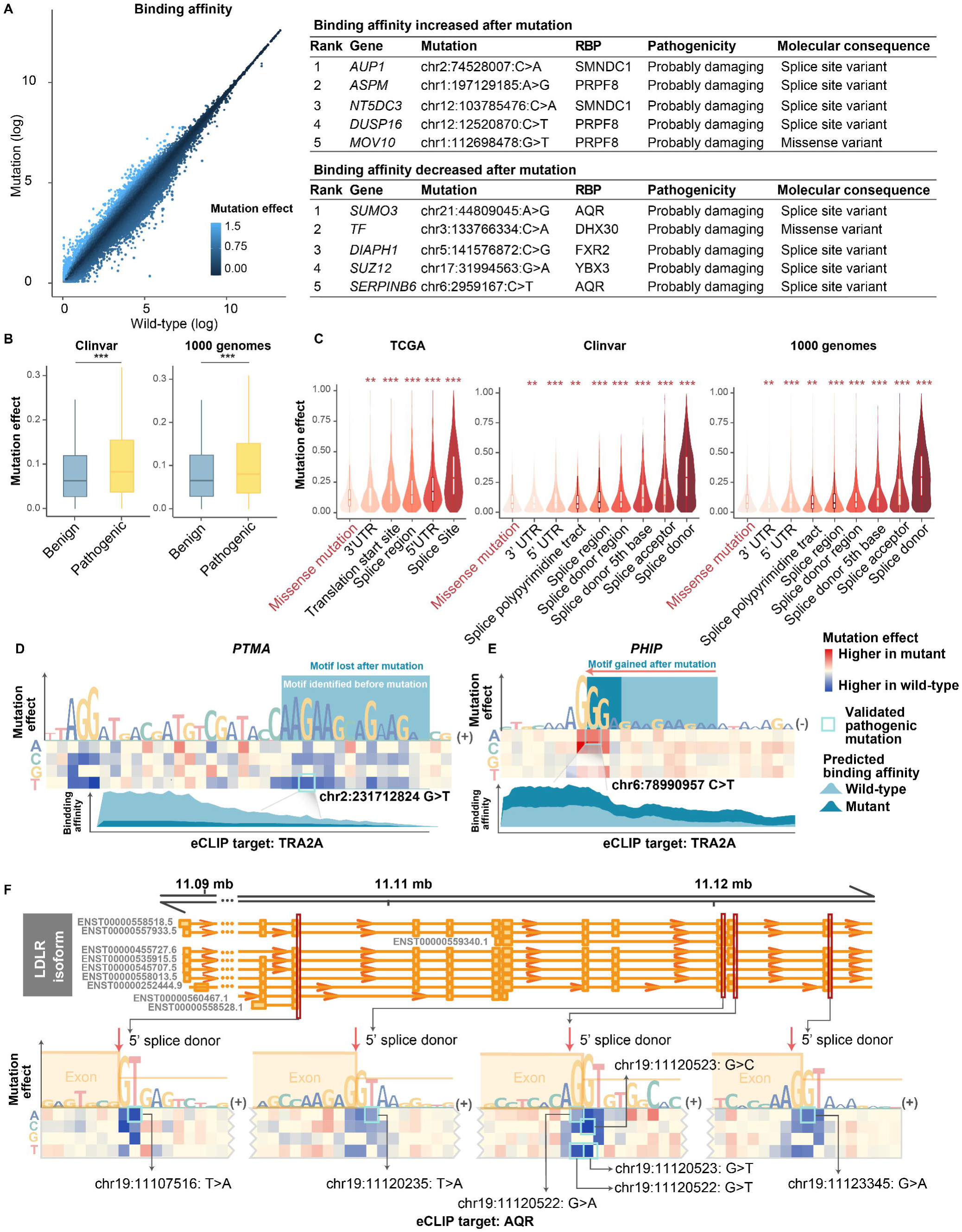
Analysis of genomic variants with RNA regulatory potentials. (**A**) Left, the predicted binding affinity (log_2_(1+x) transformed) of the wild-type (x-axis) and mutated (y-axis) sequence. Each dot represented a single nucleotide variant, with color density indicating the mutation effect score of the SNVs. Right, the top SNVs with the highest mutation effect scores. (**C**) The mutation effect scores of SNVs from distinct genome locations. The significance of the difference in mutation scores between missense mutations and splice-related mutations was assessed using the Wilcoxon test. \**p*-value < 0.05, \*\**p*-value < 0.01, \*\*\**p*-value < 0.001. (**D-F**) The representative examples of SNVs that disrupt RNA regulation. Heatmap is used to show the mutation effects of the SNVs. The known disease-associated SNVs annotated by ClinVar and the 1000 Genomes Project are highlighted with blue boxes. The color intensity in heatmap represents the mutation effect scores. The height of sequence logo plot represents the maximum mutation effect score of the position. In (**D-E**), the light blue box represents the motif positions before mutation, and the dark blue box represents the motif after mutation. The peaks reflect the predicted influence of clinically validated SNVs (highlighted with blue box in heatmap) on RBP binding affinity.

Reformer captured the impact of pathogenic mutations on RNA regulation. For example, the chr17:43106478 A>C mutation that was predicted to increase binding affinity is located at the donor site of *BRCA1* (**Supplementary Figure 9A**). This mutation was reported to promote tumorigenesis via disrupting the function of *BRCA1* [16]. Besides, we analyzed the high-scoring variations in *NF1*. Mutations in acceptor of *NF1* such as chr17:31337818 G>C and chr17:31337817 A>C was predicted to disrupt LIN28B binding. The pathogenicity of the mutations has been verified in patients with neurofibroma (**Supplementary Figure 9B**) [17,18]. One notable mutation is chr2:231712824 G>T that deletes the AAGAAGAA motif of TRA2A and reduces the binding affinity of TRA2A to RNA (**Figure 5D**). This mutation was also predicted to be deleterious by PolyPhen and SIFT (**Supplementary Table 5**) [19,20]. **Figure 5E** shows how a deleterious gain-of-function mutation creates a TRA2A binding motif in donor site of PHIP (**Supplementary Table 5**). Similarly, the deleterious mutation chr19:10681313 G>C in ILF3 cause the loss of TRA2A binding site and weaken the binding affinity of TRA2A to RNA (**Supplementary Figure 9C**). The chrX:153690374 G>A variant in SLC6A8 causes a gain of binding site for HNRNPK and increases the binding affinity to RNA (**Supplementary Figure 9D**). Specifically, LDLR and LMNA are enriched with the most high-scoring mutations prioritized by Reformer. All of the identified variations were located at RNA splicing sites (**Figure 5F** and **Supplementary Figure 9E**) and were associated with development of genetic disease [21–23].

### Experimental validation of the predicted mutations that influence RBP binding

To verify the prediction reliability of Reformer, we carried out electrophoretic mobility shift assays (EMSA) to assess whether pathogenic SNVs with high predicted mutation effect on PRPF8 occupancy truly alter RNA-protein interactions (**Figure 6A, B** and **Supplementary Table 6**). For each SNV, we synthesized paired cy3-labeled RNA probes carrying the wild type allele or mutant counterpart. An increasing amount of recombinant RNA-binding domain of PRPF8 (1760-1989 amino acid; **Supplementary Figure 10**) was incubated with a fixed amount of Cy3-labelled RNA, and the bound RNA-protein complex was resolved from the free RNA using non-denaturing gel electrophoresis. The relative abundance of free and bound RNA was then detected with chemical fluorescence imaging. Consistent with Reformer’s prediction, the mutant RNAs of *ALDH3A2* gave rise to higher binding efficiency of PRPF8 than its cognate allele in a protein dose dependent manner, and the mutant RNAs of *GPC3* give rise to lower the binding affinity than its cognate allele (*p*-value < 0.05; **Figure 6D, E** and **Supplementary Table 7**). By means of similar strategies, we confirmed that the binding of U2AF2 (150-462 amino acid; **Supplementary Figure 10**) to its RNA substrates was weakened by one pathogenic SNV with the top predicted mutation effect (**Figure 6G**, H and **Supplementary Table 7**). These findings consolidate the application of Reformer in discovering candidate SNVs that influence RBP regulation.

**Figure 6.**
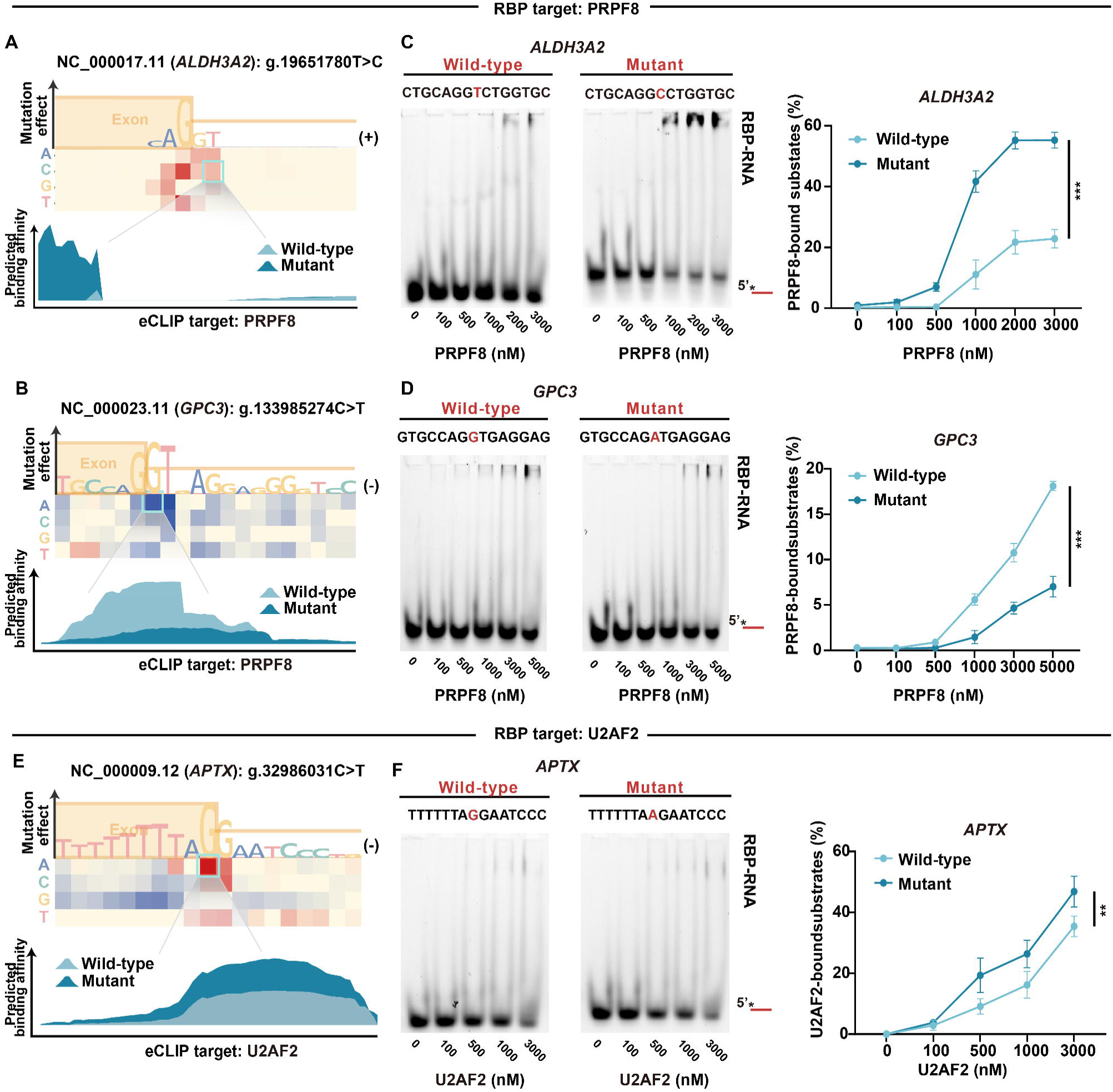
Experimental validation of SNVs altering the binding affinity to RBPs. (**A-B**) The predicted mutation effects for SNVs that alter interaction with PRPF8. Heatmap was used to show the mutation effects of the SNVs. The color intensity represents the mutation effect scores calculated with Reformer. The height of sequence logo plot represents the maximum mutation effect score of the position. The peaks reflect the predicted influence of clinical validated SNVs (highlighted with blue box in heatmap) on RBP binding affinity. (**C-D**) Left panel, electrophoretic mobility shift assays (EMSA) results of PRPF8 binding towards their RNA substrates. EMSAs were performed with increasing amount of RBPs. Right panel, of the proportion of RBPs-bound RNA. (**E**) The predicted mutation effects for SNVs that alter interaction with U2AF2. Heatmap was used to show the mutation effects of the SNVs. The color intensity represents the mutation effect scores calculated with Reformer. The height of sequence logo plot represents the maximum mutation effect score of the position. The peaks reflect the predicted influence of clinical validated SNVs (highlighted with blue box in heatmap) on RBP binding affinity. (**F**) Left panel, electrophoretic mobility shift assays (EMSA) results of U2AF2 binding towards their RNA substrates. EMSAs were performed with increasing amount of RBPs. Right panal, the proportion of RBPs-bound RNA. Three independent experiments are shown. Data are presented as mean ± s.d. \**p*-value < 0.05, \*\**p*-value < 0.01, \*\*\**p*-value < 0.001, analyzed with two-way ANOVA test.

## Discussion

Characterizing protein-RNA interactions to gain mechanistic understanding of RBP regulation is a long-standing challenge. To address this challenge, we developed Reformer to predict RNA-protein binding based solely on sequence. The attention mechanism underlying Reformer boosts the information flow across the sequence, capturing the interactions between peaks and their surrounding contexts. The effective use of sequence information leads to more accurate prediction of RBP- and cell-type-specific binding affinities at single-base resolution. Positive experimental validation results of point mutations influencing RBPs’ binding affinity support the ability of Reformer in quantifying mutations that influence RBP regulation. We anticipate that Reformer will be valuable in furthering our understanding of the RNA regulatory mechanisms underlying pathogenic mutations.

The accurate prediction of Reformer offers insights into the rule of RNA-RBP interaction. Reformer can recognize RBP binding motifs in peak regions and their surrounding contexts. For example, Reformer discovered the TTTTT motif of U2AF65 located at the left side of eCLIP-seq peaks (**Figure 3D**). This motif was confirmed to affect U2AF65 binding by controlling the conformational equilibrium of RNA [24]. Besides, we compiled reference motif signatures based on the consensus sequences identified by Reformer. The canonical motifs can be reconstructed by deconvoluting against these reference signatures, indicating that potential RBP recognition patterns are compiled in the signatures (**Figure 4B**). Linking RBP functions to motif signatures provides clues for the provenience of the signatures. For instance, Signature 32 (i.e., TAAAA) is associated with motifs of RNA-modifying proteins such as HNRNPA2B1, SRSF3, SRSF10 and A1CF (**Figure 4F**). This suggests that Signature 32 may represent a consensus binding pattern of RNA modification proteins [46]. Further investigating of the biological meanings of these motif signatures may lead to better understanding the mechanisms of RBP regulation.

In recent years, clinical research has identified numerous pathogenic mutations, yet their underlying mechanisms remain elusive [15]. Our study aims to shed light on this matter by focusing on mutations predicted to disrupt RNA binding with PRPF8, a nucleotide protein located at the catalytic core of the spliceosome [25]. Two mutations are predicted to exert the most significant influence on RNA binding to PRPF8, both of which have been previously linked to disruptive changes in protein functionality [26]. Utilizing EMSA, we experimentally confirmed that these mutations altered the binding of RNAs to PRPF8 (**Figure 6A-D**). Furthermore, our validation efforts extended to the mutations predicted to disturb RNA binding to U2AF2, a critical pre-mRNA splicing factor [27]. Particularly, the mutation predicted with the greatest influence on U2AF2 binding affinity has been confirmed to be associated with Ataxia-oculomotor apraxia syndrome (rs1563963464). Our EMSA-based validation demonstrated the change of binding between U2AF2 and RNA caused by this mutation (**Figure 6E-F**). These findings provide evidence for understanding the pathogenic mechanisms associated with these pathogenic mutations.

Reformer has several advantages. First, Reformer purely takes the sequence as input for prediction, which makes the model input more readily available than methods relying on both sequence and structure information [8,44]. As more experimental data becomes available [1], Reformer is promising for characterizing RNA-protein interactions over a broader range of RBPs and cellular conditions. Second, Reformer tasks with binding affinity prediction at base resolution, which is more precise than previous methods for distinguishing binding from non-binding sites [6,8,45]. Third, Reformer fits all eCLIP-seq targets with a single model, effectively reducing the consumption of model training as compared with previous models [8,45]. Last, while previous models focus on peak regions of about 100 bp [6,8,45], Reformer expands the receptive field to 512 bp, allowing the model to learn contextual information. Altogether, these advantages accelerate the motif mining and pathogenic variant discovery.

Reformer can be boosted in several aspects. Firstly, enhancing the quality of experimental data [1], and lengthening the receptive region to allow for long-range contextual learning [12], would likely improve performance. Secondly, the model can only model RNA-protein interactions in limited RBPs and biological conditions currently. The growing amount of experimental data will expand its compatibility. Finally, the sensitivity of the model to genetic mutations could be further improved by fine-tuning upon well-annotated mutation datasets.

We anticipate that Reformer will be useful in discovering motifs, and mapping pathogenic mutations to their effects on RBP regulation. We have made the pre-trained Reformer publicly available to boost these downstream applications. We hope that Reformer will contribute to a better understanding of the regulatory mechanisms that govern RBP regulation and pave new ways to improve diagnosis and therapeutic modalities for diseases.

## Methods

### RBP binding data collection and processing

We collected 225 eCLIP-seq experiments from the encyclopedia of DNA elements (ENCODE) repository [28] before March 2021. The eCLIP-seq targets include 155 RBPs across 3 cell lines (**Supplementary Table 1**). Experimental replicates of eCLIP-seq were collected from ENCODE repository up to October 2022.

We defined peaks as regions with a log_2_(fold-enrichment coverage) of at least 3 and a –log_10_(*p*-value) of at least 5 as compared with a size-matched control [29]. The peaks ranged from 1 to 426 bp in length, which we expanded to 511 bp from the middle to both sides. We then extracted the corresponding coverage of the eCLIP-seq peaks and mapped the peaks to GRCh38.p5 sequences. We randomly divided the chromosomes into training, validation and test set, containing 872,618, 23,633 and 94,713 sequences, respectively (**Supplementary Table 1**). To normalize the peak coverages for each eCLIP-seq experiment, we divided the absolute coverage by the sum of all coverages, and set peak coverages above 2500 to 2500.

### Model architecture

We used a bidirectional encoder representation from transformers [30] for RNA-protein binding learning. The model comprised three main components: (1) a sequence encoder layer; (2) 12 transformer layers; (3) a linear layer for predicting binding affinity. The model took as input the cDNA sequence with a max length of 511 bp, and predicted binding affinity at base resolution as output. To tokenize each sequence, we represented it with a 3-mer representation and add the corresponding eCLIP-seq target name as one token before the sequence. We further included a special token [CLS] at the beginning and a special token [SEP] at the end of the sequence. For example, the DNA sequence ‘ATCGA’ from SRSF1 target of K562 cell line was represented with six tokens: {[CLS], SRSF1&K562, ATC, TCG, CGA, [SEP]}. We represented the sequence into a matrix *M* by embedding each token into a numerical vector. Additionally, we incorporated position information by adding absolute position embedding in the network. The position information was embedded as trainable parameters, which were randomly initialized and updated during training. The token embedding and position embedding were summarized and input into the transformer layers.

The Reformer model consists of 12 transformer layers, each with 12 attention heads and 768 hidden units for protein-RBP interaction learning. The transformer layer captures token-token interaction information using a multi-head self-attention mechanism [31]. Given an input, the self-attention mechanism assigns an attention weight α *_i,j_* > 0 to each token pair *i,j*, where ∑ *_j_* α *_i,j_* = 1. Attention in the transformer is bidirectional. Attention weights α _i,j_ are computed by the scaled dot-product of the query vector of *i* (*Q*) and the key vector of *j* (*K*), followed by a softmax operation. Then attention weights are used to produce a weighted sum of value vectors (*V*):

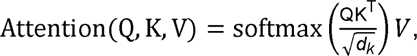

where *d _k_* is the dimension of *K*. In the multi-layers, multi-heads setting, the attention scores were different in each layer and head.

The output of the last transformer layer was input into a linear layer for predicting binding affinity. The linear layer predicted binding coverage for each of the 509 bp sequence and trims the last 2 bases on the right side, which was not fully tokenized. The predicted values were then passed to a rectified linear unit (ReLU) activation function, which returns zero for negative input and the input itself for non-negative input:

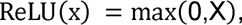

we set the dropout rate of 0.1 for both the transformer layers and the linear layer. A detailed model architecture was shown in **Supplementary Figure 1**.

### Model training and evaluation

The model was initialized with the parameters of a pre-trained model [32]. We fine-tuned the model to minimize the mean-squared loss function, which was defined as the mean squared error between the reference (*Y*) and predicted coverage (*I*):

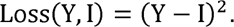

The model was trained at a learning rate of 2e-5 for 30 epochs and the mini-batch stochastic gradient descent algorithm [33], with a weight decay of 1e-4. The model was implemented using PyTorch (version 1.7.1) and transformers (version 4.10.0), and was run on an NVIDIA DGX A100 system with 8 GPUs, each with 40 Gb of memory.

We utilized the validation set to select hyper-parameter and the test set to evaluate the model performance. We considered two evaluation metrics to assess the model performance: (1) at base resolution, we calculated the Pearson correlation between the log_10_(1+x) transformed measured and predicted base coverage; (2) at sequence level, we calculated the Pearson correlation between the measured and predicted peak binding affinity, which was defined as the sum of log_10_(1+x) transformed base coverage across the peak; (3) we also measured the coverage differences between the predicted and measured peaks, which was represented as:

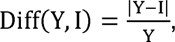

where *Y* and *I* represented the log_10_(1+x) transformed measured and predicted binding affinity, respectively.

### Motif enrichment analysis in regions that highly attended by Reformer

For each sequence, we extracted the attention between bases as a 512 * 512 matrix per attention head (*F _i, j_*, where *i* and *j* = 1, 2, …, 512 corresponding to each base of the sequence). We then applied average product correlation (APC) regression on the attention matrix to remove noise [34]. Specifically, APC regression was computed as:

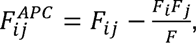

where *F _i_*, *F _j_*, and *F* represent the sum of attention over *i*-th row, *j*-th column, and full matrix, respectively.

We defined the binding sites as peak regions with binding affinity > 10, and calculated the attention score between the binding sites with other sites. For each base, the attention score was represented as the attention summation across itself to all binding sites.

To identify regions that were highly attended by Reformer, we employed a sliding window approach, where each sequence was divided into windows of size 10 bp with a step size of 1 bp. We then summed up the attention scores for each window and defined regions that were highly attended by Reformer as the top 1% of sliding windows with the highest attention scores. Two regions were iteratively merged if they have at least 1 bp overlap.

We validated whether the highly attended regions were enriched for motifs using AME algorithm [35]. For each eCLIP-seq target, we collected motifs of the corresponding RBP from ATract database [36]. A total of 920 motifs less than 10 bp were curated (**Supplementary Table 2**). The controls were randomly selected from regions that were not highly attended by Reformer. The motif enrichment analysis was performed for each motif per attention head. As a comparative analysis, we performed motif enrichment analysis in eCLIP-seq peak regions. The non-peak regions in 511 bp across the peaks were selected as controls. A Fisher’s exact test *p*-value < 0.05 was considered significant.

### Motif signature construction

For each attention head, we extracted the top 1% windows the highest attention scores based on the sliding window strategy described above. The window size was set to 5 bp and the window step was set to 1 bp. We then summed up the attention scores of each alphabet at each position and calculated their cross-entropy, resulting in a position weight matrix (PWM) of 4 * 5 matrix (*p _i, j_* where *i* = 1, 2, 3, 4 corresponding to A, C, G, T, and *j* is site index, 1 ≤ *j* ≤ 5) as the motif signature. *P*-values were calculated with permutation test by computing a test statistic on the motif signature and then for 1000 permutations of the random generated motifs.

We collected 1312 motifs of ATtract database and reconstructed them against the motif signatures using nonnegative matrix factor 2-D deconvolution (NMF2D) [37] algorithm. The NMF2D algorithm was formulated as follows:

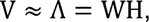

where the non-negative *V,* D*, W* and *H* represented the real motif matrix, the reconstructed motif matrix, the motif signature matrix and the weight matrix, respectively. The goal of NMF2D is to minimize a generalized Kullback-Leibler divergence to optimize *H* for reconstructing *V*:

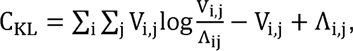

*V _i,j_* and □ *_i,j_* denote the real and reconstructed PWM of *j*-th base in *i*-th motif, respectively. *P*-values were calculated with permutation test by computing a test statistic on the motif signature and then for many permutations of the random generated signatures.

The similarity between the real motif matrix *V* and the reconstructed motif matrix was measured with TOMTOM motif similarity algorithm [35], and motifs with a *q*-value < 0.05 was considered significant. We validated the motif reconstruction performance for motif signatures ranging from 5 to 10 bp (**Supplementary Table 4**). We present the motif signatures of 5 bp in the manuscript.

To investigate whether the motif signatures were associated with specific RNA processing functions, we used the weight matrix *H* to quantify the contribution of the signatures to motifs. We collected domain and functional annotations of RBPs from the previous studies [38–44], and the corresponding motifs were collected from ATract database [36]. Statistical analysis was performed using the Wilcoxon rank-sum test, with a significance threshold of a Benjamini-Hochberg adjusted *p*-value < 0.05. We displayed the motifs with the highest contributions in **Figure 4D**.

### Benchmarking mutation effect predictions on saturation mutation data

Disease-related variants were obtained from Clinvar database [14], the 1000 Genomes Project [15] and the TCGA repository in November 2022. We extracted the single nucleotide variants for downstream analysis. We select variants with clinical significance as pathogenic or benign and excluded the variants that had conflicting clinical significance from the ClinVar database and the 1000 Genomes Project. A total of 73,022, 84,208 and 396,573 SNVs with curated annotations were retrieved from ClinVar [14], the 1000 Genomes Project [15] and the TCGA repository, respectively. The genome locations of splice sites were extracted from standard genome annotation GENCODE [45] (release 24 of GRCh38). For SNVs in 100 Genomes Project, common variants were defined as mutations with an allele frequency >1%, while the rare variants were defined as mutations with an allele frequency <0.1%. We overlapped the SNP locus with eCLIP-seq coverage >10 for mutation effect calculation.

For each variant, we evaluated its mutation effect as the changes in predicted binding affinity before and after mutation:

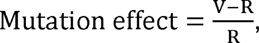

the binding affinity was measured as the coverage summation of 100 bp around the mutated nucleotide, denoted by R and V before and after mutation, respectively.

We validated the pathogenic mutations that were predicted to disrupt RNA binding to PRPF8. We specifically focused on mutations from the 1000 Genomes Project that had binding coverage of mutant or wild-type RNA to PRPF8 exceeding 150. We selected the top 1 and 3 mutations that displayed the highest predicted influence on PRPF8 binding affinity for experimental validation. Employing the same selection process, we further experimental validated the foremost pathogenic mutation predicted to perturb RNA binding to U2AF2.

### Purification of recombinant human PRPF8 and U2AF2

The cDNA fragments encoding the PRPF8 1760-1989 amino acid and U2AF2 150-462 amino acid were cloned into pET28a vector. The human PRPF8-pET28a and U2AF2--pET28a expression vectors were transformed into BL21 (DE3) chemically competent cells (TransGen, CD601-02). Protein induction was carried out via 0.1 mM isopropyl β-D-1-thiogalactopyranoside (IPTG) treatment and cells were lysed with sonication in ice-cold buffer 1 [25 mM Tris-HCl (pH7.5), 500 mM NaCl, 5mM imidazole (pH7.5), 3 mM 2-Mercaptoethanol, 0.1 mM phenylmethanesulfonylfluoride or phenylmethylsulfonyl fluoride (PMSF)]. Subsequently, the lysate was centrifuged at 15,000 rpm for 45 min at 4 ℃. The supernatant was loaded onto the HIS-Select Nickel Affinity Gel (sigma, P6611) followed by washing the beads with buffer 1. The sample was eluted with buffer 2 [25 mM Tris-HCl (pH7.5), 2000 mM NaCl, 3 mM 2-Mercaptoethanol]. The fusion protein was eluted with buffer 4 [25 mM Tris-HCl (pH7.5), 500 mM NaCl, 300 mM imidazole (pH7.5), 3 mM 2-Mercaptoethanol]. The concentrated protein was then loaded into the HiTrap Heparin HP column (Cytiva, 17-0407-01) and eluted with a linear gradient of 0.05 to 1.0 M NaCl. The eluted protein was concentrated by ultrafiltration and determined by Coomassie blue staining.

### Electrophoretic Mobility Shift Assay

DNA substrates labelled with Cy3 (Invitrogen) at 5’-end (50 nM) were incubated with indicated amounts of proteins in 1× binding buffer [25 mM Tris (pH 7.5), 50 mM NaCl, 5 mM MgCl2, 1 mM DTT, 5% glycerol, and 0.05% Triton X-100] at room temperature for 30 min. The reaction mixture (20μL in total) was then loaded with 2μL 10 × loading dye and resolved on 8% native acrylamide/Bis gel in cold 0.5 × TBE buffer (44.5 mM Tris, 44.5 mM boric acid, and 0.5 mM EDTA, pH8.3). Signals were detected with an Alliance Q9 imager, and band intensities were quantified by ImageJ (NIH).

## Code availability

The source code of Reformer is available at https://github.com/xilinshen/Reformer.

## Data availability

The eCLIP-seq experiments from the encyclopedia of DNA elements (ENCODE) repository (https://www.encodeproject.org/). The motifs of RBPs are available from ATract database (http://attract.cnic.es). Disease-related variants were obtained from Clinvar database (https://www.ncbi.nlm.nih.gov/clinvar/) [14], the 1000 Genomes Project (https://www.internationalgenome.org/) [15] and the TCGA repository (https://portal.gdc.cancer.gov/). The predicted mutation effect scores for all SNPs are available at https://github.com/xilinshen/Reformer.

## Acknowledgment

This work was supported by the National Natural Science Foundation of China [32270688 and 31801117]; the Program for Changjiang Scholars and Innovative Research Team in University in China [IRT_14R40]; the Tianjin Science and Technology Committee Foundation [17JCYBJC25300]; and the Chinese National Key Research and Development Project [2018YFC1315600]. We want to thank all the researchers for their generosity to make their data publicly available.

## Supplementary figure and table legends

**Supplementary Figure 1. The model architecture of Reformer.** The model was constructed with a sequence encoder layer, 12 transformer blocks and a regression layer for binding affinity prediction. The layer shapes (without batch dimensions) are shown as tuples followed the layer names.

**Supplementary Figure 10.** Coomassie brilliant blue staining of His-tagged U2AF2 and PRPF8 purified from bacterial cells.

**Supplementary Table 1.** Description of 255 eCLIP-seq experiments and 5 replications used for model training and evaluation.

**Supplementary Table 2.** Motif enrichment results of Reformer and eCLIP-seq experiments.

**Supplementary Table 3.** The statistical significance of the motif signatures.

**Supplementary Table 4.** Detailed information of the motifs reconstructed with motif signatures.

**Supplementary Table 5.** The 1500 SNVs with the highest mutation effect scores in ClinVar, TCGA and the 1000 Genomes Project.

**Supplementary Table 6.** The predicted mutation effects for SNVs with experimental validation.

**Supplementary Table 7.** Electrophoretic mobility shift assays (EMSA) results of RBPs binding towards their RNA substrates.

## References

1. Van Nostrand EL, Freese P, Pratt GA, et al. A large-scale binding and functional map of human RNA-binding proteins. Nature 2020; 583:711–719

2. Montes M, Sanford BL, Comiskey DF, et al. RNA Splicing and Disease: Animal Models to Therapies. Trends Genet. 2019; 35:68–87

3. Sade-Feldman M, Yizhak K, Bjorgaard SL, et al. Defining T Cell States Associated with Response to Checkpoint Immunotherapy in Melanoma. Cell 2018; 175:998–1013.e20

4. Ferlini A, Goyenvalle A, Muntoni F. RNA-targeted drugs for neuromuscular diseases. Science (80-.). 2021; 371:29–31

5. Shuai S, Suzuki H, Diaz-Navarro A, et al. The U1 spliceosomal RNA is recurrently mutated in multiple cancers. Nature 2019; 574:712–716

6. Alipanahi B, Delong A, Weirauch MT, et al. Predicting the sequence specificities of DNA- and RNA-binding proteins by deep learning. Nat. Biotechnol. 2015; 33:831–838

7. Uhl M, Tran VD, Heyl F, et al. RNAProt: an efficient and feature-rich RNA binding protein binding site predictor. Gigascience 2021; 10:

8. Sun L, Xu K, Huang W, et al. Predicting dynamic cellular protein–RNA interactions by deep learning using in vivo RNA structures. Cell Res. 2021; 31:495–516

9. Taliaferro JM, Lambert NJ, Sudmant PH, et al. RNA Sequence Context Effects Measured In Vitro Predict In Vivo Protein Binding and Regulation. Mol. Cell 2016; 64:294–306

10. Vaswani A, Shazeer N, Parmar N, et al. Attention Is All You Need. Arxiv 2017;

11. Devlin J, Chang M-W, Lee K, et al. BERT: Pre-training of Deep Bidirectional Transformers for Language Understanding. Arxiv 2019;

12. Avsec Ž, Agarwal V, Visentin D, et al. Effective gene expression prediction from sequence by integrating long-range interactions. Nat. Methods 2021; 18:1196–1203

13. Gerber AP, Herschlag D, Brown PO. Extensive Association of Functionally and Cytotopically Related mRNAs with Puf Family RNA-Binding Proteins in Yeast. PLoS Biol. 2004; 2:e79

14. Landrum MJ, Lee JM, Benson M, et al. ClinVar: improving access to variant interpretations and supporting evidence. Nucleic Acids Res. 2018; 46:D1062–D1067

15. Auton A, Abecasis GR, Altshuler DM, et al. A global reference for human genetic variation. Nature 2015; 526:68–74

16. Cochran RL, Cidado J, Kim M, et al. Functional isogenic modeling of BRCA1 alleles reveals distinct carrier phenotypes. Oncotarget 2015; 6:25240–25251

17. Wilson BN, John AM, Handler MZ, et al. Neurofibromatosis type 1: New developments in genetics and treatment. J. Am. Acad. Dermatol. 2021; 84:1667–1676

18. Baralle D. Splicing in action: assessing disease causing sequence changes. J. Med. Genet. 2005; 42:737–748

19. Adzhubei I, Jordan DM, Sunyaev SR. Predicting Functional Effect of Human Missense Mutations Using PolyPhen-2. Curr. Protoc. Hum. Genet. 2013; 76:

20. Ng PC. SIFT: predicting amino acid changes that affect protein function. Nucleic Acids Res. 2003; 31:3812–3814

21. Takada D, Emi M, Ezura Y, et al. Interaction between the LDL-receptor gene bearing a novel mutation and a variant in the apolipoprotein A-II promoter: molecular study in a 1135-member familial hypercholesterolemia kindred. J. Hum. Genet. 2002; 47:0656–0664

22. Mozas P, Castillo S, Tejedor D, et al. Molecular characterization of familial hypercholesterolemia in Spain: Identification of 39 novel and 77 recurrent mutations in LDLR. Hum. Mutat. 2004; 24:187–187

23. Lee JM, Nobumori C, Tu Y, et al. Modulation of LMNA splicing as a strategy to treat prelamin A diseases. J. Clin. Invest. 2016; 126:1592–1602

24. Mackereth CD, Madl T, Bonnal S, et al. Multi-domain conformational selection underlies pre-mRNA splicing regulation by U2AF. Nature 2011; 475:408–411

25. Grainger RJ, Beggs JD. Prp8 protein: At the heart of the spliceosome. RNA 2005; 11:533–557

26. Kraus C, Braun-Quentin C, Ballhausen W, et al. RNA-based mutation screening in German families with Sjögren-Larsson syndrome. Eur. J. Hum. Genet. 2000; 8:299–306

27. Sutandy FXR, Ebersberger S, Huang L, et al. In vitro iCLIP-based modeling uncovers how the splicing factor U2AF2 relies on regulation by cofactors. Genome Res. 2018; 28:699–713

28. . An integrated encyclopedia of DNA elements in the human genome. Nature 2012; 489:57–74

29. Van Nostrand EL, Pratt GA, Shishkin AA, et al. Robust transcriptome-wide discovery of RNA-binding protein binding sites with enhanced CLIP (eCLIP). Nat. Methods 2016; 13:508–514

30. Devlin J, Chang M-W, Lee K, et al. BERT: Pre-training of Deep Bidirectional Transformers for Language Understanding. 2018;

31. Shaw P, Uszkoreit J, Vaswani A. Self-Attention with Relative Position Representations. 2018;

32. Ji Y, Zhou Z, Liu H, et al. DNABERT: pre-trained Bidirectional Encoder Representations from Transformers model for DNA-language in genome. Bioinformatics 2021; 37:2112–2120

33. Li M, Zhang T, Chen Y, et al. Efficient mini-batch training for stochastic optimization. Proc. 20th ACM SIGKDD Int. Conf. Knowl. Discov. data Min. 2014; 661–670

34. Dunn SD, Wahl LM, Gloor GB. Mutual information without the influence of phylogeny or entropy dramatically improves residue contact prediction. Bioinformatics 2008; 24:333–340

35. Bailey TL, Johnson J, Grant CE, et al. The MEME Suite. Nucleic Acids Res. 2015; 43:W39–W49

36. Giudice G, Sánchez-Cabo F, Torroja C, et al. ATtRACT—a database of RNA-binding proteins and associated motifs. Database 2016; 2016:baw035

37. Schmidt MN, Mørup M. Nonnegative Matrix Factor 2-D Deconvolution for Blind Single Channel Source Separation. 2006; 700–707

38. Smialek MJ, Ilaslan E, Sajek MP, et al. Role of PUM RNA-Binding Proteins in Cancer. Cancers (Basel). 2021; 13:129

39. Zhang A, Liu W-F, Yan Y-B. Role of the RRM domain in the activity, structure and stability of poly(A)-specific ribonuclease. Arch. Biochem. Biophys. 2007; 461:255–262

40. Olejniczak M, Jiang X, Basczok MM, et al. KH domain proteins: Another family of bacterial RNA matchmakers? Mol. Microbiol. 2022; 117:10–19

41. Larsen NA. The SF3b Complex is an Integral Component of the Spliceosome and Targeted by Natural Product-Based Inhibitors. 2021; 409–432

42. Rino J, Desterro JMP, Pacheco TR, et al. Splicing Factors SF1 and U2AF Associate in Extraspliceosomal Complexes. Mol. Cell. Biol. 2008; 28:3045–3057

43. Lewis CJT, Pan T, Kalsotra A. RNA modifications and structures cooperate to guide RNA–protein interactions. Nat. Rev. Mol. Cell Biol. 2017; 18:202–210

44. Naganuma T, Nakagawa S, Tanigawa A, et al. Alternative 3′-end processing of long noncoding RNA initiates construction of nuclear paraspeckles. EMBO J. 2012; 31:4020–4034

45. Frankish A, Diekhans M, Ferreira A-M, et al. GENCODE reference annotation for the human and mouse genomes. Nucleic Acids Res. 2019; 47:D766–D773

